# CRISPR screening of porcine sgRNA library identified host factors essential for Japanese encephalitis virus replication

**DOI:** 10.1101/840835

**Authors:** Changzhi Zhao, Hailong Liu, Tianhe Xiao, Zichang Wang, Xiongwei Nie, Xinyun Li, Ping Qian, Liuxing Qin, Xiaosong Han, Jinfu Zhang, Jinxue Ruan, Mengjin Zhu, Yiliang Miao, Bo Zuo, Kui Yang, Shengsong Xie, Shuhong Zhao

**Author notes:** To whom correspondence should be addressed. Shuhong Zhao, Huazhong Agricultural University, No.1, Shizishan Street, Hongshan District, Wuhan, Hubei Province, 430070, P. R. China.; Correspondence may also be addressed to Shengsong Xie, Huazhong Agricultural University, No.1, Shizishan Street, Hongshan District, Wuhan, Hubei Province, 430070, P. R. China. The authors contributed equally (Changzhi Zhao, Hailong Liu, Tianhe Xiao).

## Abstract

Japanese encephalitis virus (JEV) is a mosquito-borne zoonotic flavivirus that causes encephalitis and reproductive disorders in mammalian species. However, key host genes involved in the JEV life cycle and cell death are poorly understood. Here, we designed 85,674 single guide RNAs (sgRNAs) targeting 17,743 protein-coding genes, 11,053 long ncRNAs (lncRNAs), and 551 microRNAs (miRNAs) in the porcine genome, and subsequently developed a porcine sgRNA library and genome-scale CRISPR/Cas9 knockout (PigGeCKO) system. These sgRNAs were delivered into porcine kidney-15 (PK-15) cells that constitutively express Cas9, positive selection screening of the resulting PigGeCKO cell collection for resistance to JEV-induced cell death led to the identification of several previously unreported genes required for JEV infection. We conducted follow-up studies to verify the dependency of JEV on these genes, and identified functional contributions for six of the many candidate JEV-related host genes, including an endoplasmic reticulum membrane protein complex subunit 3 (EMC3) and calreticulin (CALR). Additionally, we identified that four genes associated with heparan sulfate proteoglycans (HSPGs) metabolism, specifically those responsible for HSPG sulfurylation, facilitated JEV entry into PK-15 cells. Thus, beyond our development of the largest CRISPR-based functional genomic screening platform for pig research to date, this work deepens our basic understanding of flavivirus infection and identifies multiple potentially vulnerable targets for the development of medical and breeding technologies to prevent and treat diseases caused by Japanese encephalitis virus.

## INTRODUCTION

Numerous viruses in nature are capable of infecting both humans and domestic animals. Among these is Japanese encephalitis virus (JEV), a flavivirus which is closely related to Dengue Virus (DENV), Zika Virus (ZIKV), Yellow Fever Virus (YFV), West Nile Virus (WNV) and Hepatitis C Virus (HCV)^1,2^. JEV is the leading cause of viral encephalitis in humans in some Asian countries, with an estimated 69,000 severe clinical cases and approximately 13,600 to 20,400 deaths annually^3^, despite the widespread use of vaccine. While several inactivated and live vaccines are used to prevent JEV infection, no antiviral drugs are available for the treatment of JEV-related diseases^4,5^. Despite achievements in control and prevention of JEV infection, this disease remains a major public health concern in Northern Oceania and in South, East, and Southeast Asia, and is viewed as an emerging global pathogen^6^.

JEV is a mosquito-borne virus that can be greatly amplified in pigs, causing swine encephalitis and reproductive complications in this species. Pigs are readily infected with JEV and can develop high levels of viremia^7,8^. JEV infection is usually asymptomatic in adult pigs, but manifestations of abortion, still-birth, and birth defects, including central nervous system defects, are not unusual following infection in pregnant swine, which can result in substantial economic losses to pork producers^9,10^. Accordingly, JEV outbreaks represent a major threat to both public health and the agricultural economy, especially in areas with low vaccine coverage and/or limited diagnostic capacity^11^. Currently, JEV is only endemic in the Asia-Pacific region, however it was previously shown that domesticated pigs could potentially become amplification hosts upon the introduction of JEV in other countries^12^.

The JEV infection cycle starts with binding to unknown cellular receptors and attachment factors^6,13^, followed by viral entry to enable replication. Subsequently, the JEV RNA genome is replicated, viral particles are matured and packaged, and released from cells. JEV infects a variety of cell types from diverse species (including mammals, birds, amphibians, and insects), suggesting that JEV can likely access multiple cell types using multiple host receptors^14^. In recent years, tremendous progress has been made in understanding the viral components required for the various steps of JEV entry and replication, but little is known about the host cell components involved in this process^15–18^. Understanding virus–host interactions by elucidating the molecular mechanisms of viral transmission can help identify potential antiviral targets for developing both prophylactic and therapeutic medicines.

Efforts to treat and prevent viral infectious have traditionally been aided by genetic screening research to improve our understanding of viral dependencies and to identify potential antiviral strategies. The emergence of CRISPR genetic screening tools has spurred a new era of efficient, versatile, and large-scale screening efforts, with notable examples for flaviviridae family viruses^19–21^. Indeed, work in human cell lines based on CRISPR-based screening strategies (with virus-induced cell death readout phenotypes) have successfully identified required host genes for infection by DENV, ZIKV, WNV, YFV, and HCV^20–24^. These studies have repeatedly illustrated that genome-scale CRISPR screening represents an extremely powerful tool for both basic biology and medical research. However, we are unaware of any genome-scale efforts to examine JEV infection; moreover, there are no reports of genome-scale CRISPR/Cas9 libraries for screening studies in pigs.

Aiming to develop such a resource, and specifically seeking to study the genetic basis of resistance against JEV infection, we developed a resource which we term PigGeCKO (for porcine genome-scale CRISPR knockout), which is comprised of a library of ~85,000 sgRNA constructs (both as plasmids and as prepared lentiviruses). After developing this genome-scale sgRNA library, we infected JEV-susceptible porcine kidney-15 (PK-15) cells^25^ stably expressing the Cas9 protein with the pooled lentiviral sgRNA library, and used Fluorescence-Activated Cell Sorting (FACS) to isolate and enrich cells harboring sgRNA constructs. We then performed positive selection screening by exposing the PigGeCKO cell collection to repeated rounds of JEV challenge, retaining the viable (i.e., JEV-resistant) cells from each round. PCR amplification and deep sequencing enabled us to detect enrichment among the candidate JEV-infection-associated genes for annotations relating to heparan sulfate proteoglycans (HSPGs) and endoplasmic reticulum-associated protein degradation (ERAD) pathways. We then generated single gene knockout cell lines for six of the candidate genes, and successfully confirmed their requirement for JEV infection of PK-15 cells. These newly discovered host genes are potential targets for the development of therapies for the treatment of Japanese encephalitis and porcine diseases caused by JEV, and can also be used in the construction of genetically edited disease-resistant animal models. We anticipate that our benchmark-setting CRISPR/Cas9 screening resources will greatly facilitate basic and applied functional genomics research in pigs.

## RESULTS

### Strategy for identifying genes essential for JEV-induced cell death in pigs

The overall development process of the PigGeCKO resources is depicted in the schematic diagram in Fig. 1a, and proceeded as follows. We initially used CRISPR-offinder (v1.2)^26^ to design 85,674 specific and predicted high-efficiency sgRNAs, that collectively targeted 17,743 protein-coding genes, 11,053 lncRNAs, and 551 miRNAs in the porcine genome, as well as 1,000 negative control sgRNA constructs predicted not to target any porcine genome loci (Fig. 1b, Supplementary Table 1). Three sgRNA constructs were designed for each targeted locus, all loci which met the selection criteria detailed in the Materials and Methods were targeted. These designed sgRNA constructs were synthesized as an oligo array, which was employed as the template for PCR amplification of the sgRNA oligos that were subsequently cloned into lentiviral vectors using Gibson assembly.

**Fig 1.**
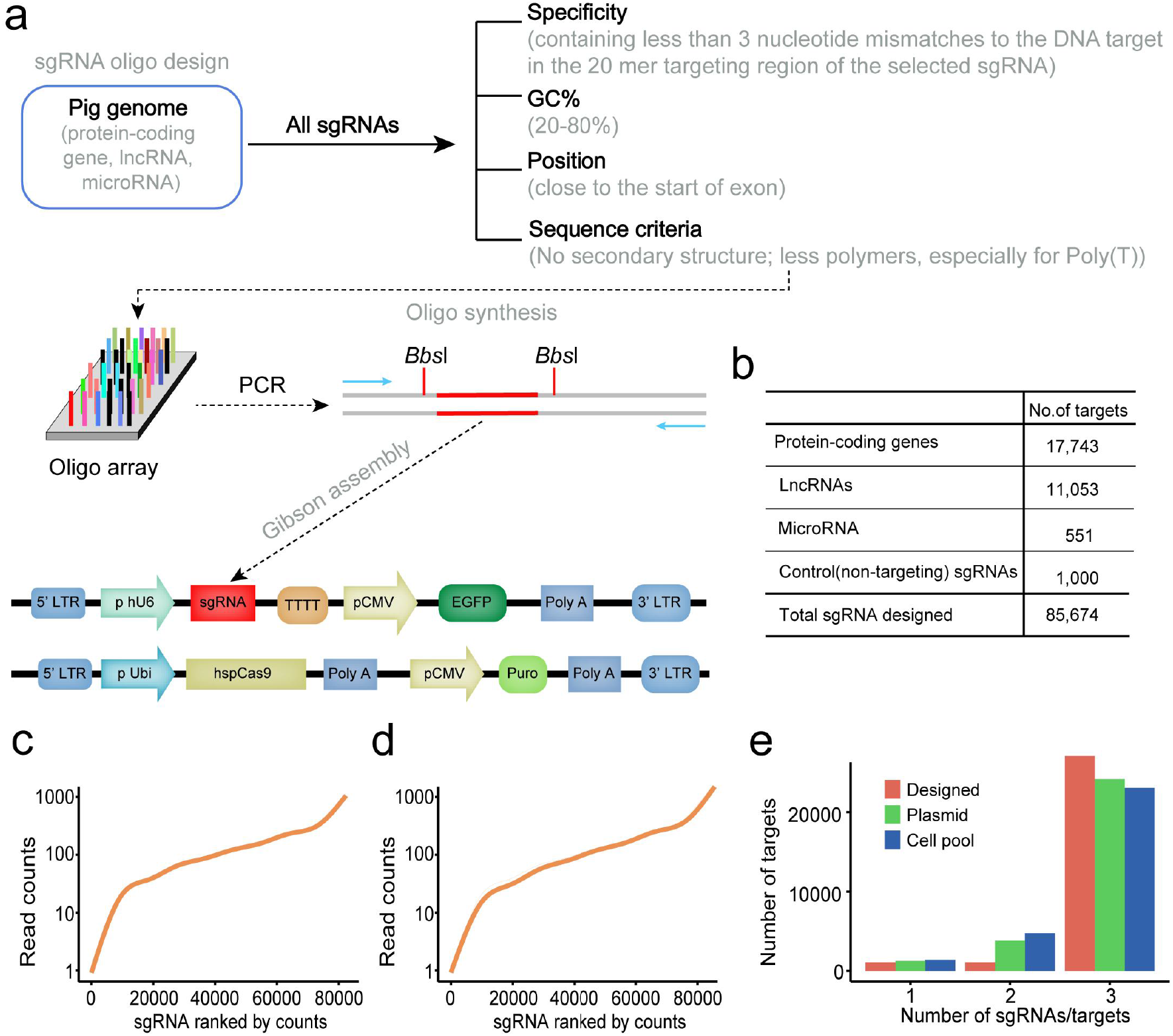
Generation of a porcine genome-wide lentiviral sgRNA library. **a**, Pipeline for sgRNA library design and construction. Protein-coding genes from Ensemble, lncRNAs from the ALDB database, and microRNAs from the miRBase database. Primers indicated by arrows were used for amplification of the sgRNA targeting sequences from the synthesized oligo array, which were cloned into the lentiviral plasmid. **b**, The number of designed sgRNA targets. **c**and **d**, Sequencing result of sgRNAs targeting sequences in the two CRISPR/sgRNA libraries. Plasmid pools (**c**) and sorted mutant cell populations (**d**) containing the whole sgRNA library were characterized using next-generation sequencing (NGS). The curve indicates the distribution of sgRNAs. **e**, The number of sgRNAs per gene in the genome-wide library from the designed, plasmid, or mutant cell pools. sgRNA: small guide RNA; LncRNA: Long non-coding RNA; Designed: sgRNA designed by CRISPR-offinder software; Plasmid: the sequencing result of sgRNA library from plasmid pools; Cell pool: the sequencing result of sgRNA library from sorted mutant cell populations.

To test the quality of the PigGeCKO plasmid library, we PCR amplified the cloned sgRNA constructs, performed deep sequencing, and found that 96.2% (82,426/85,674) of the initially designed and synthesized sgRNA sequences were present in the plasmid library (Fig. 1c, Supplementary Table 2). Although a small fraction of sgRNA were under- or over-represented, approximately 90% of the sgRNAs were within a range covering a tenfold difference in frequency (Fig. 1c). In parallel with our sgRNA library preparation, we developed a PK-15 cell line that stably expressed high levels of Cas9 (PK-15-Cas9, Clone-#14, Supplementary Fig. 1). We used an sgRNA lentivirus targeting the randomly selected *ANPEP* gene to assess the capacity of this cell line for gene editing (Supplementary Fig. 2a), and found that gene-editing activity tended to be stable approximately 6-10 days post-infection of the sgRNA-harboring lentivirus in PK-15-Cas9 cells (Supplementary Fig. 2b, c).

We next generated the PigGeCKO lentivirus library by transfecting HEK293T cells with the lentiviral sgRNA plasmids together with helper plasmids. To minimize the chance of inserting multiple sgRNAs into the same PK-15-Cas9 cells, we employed a low multiplicity of infection (MOI) to obtain a transduction rate of around 30% according to a previous study^27^. The lentivirus sgRNA library was subsequently transducted into PK-15-Cas9 cells. We performed FACS-based sorting on the signal from the green fluorescent protein (GFP) reporter which was included in all PigGeCKO constructs, infected cells with the sgRNA construct sequences were PCR amplified followed by deep sequencing. Strikingly, 94.7% (81,095/85,674) of the originally designed sgRNA sequences were retained in the PigGeCKO knockout cell collection (Fig. 1d, Supplementary Table 2). Furthermore, the plasmid library and PigGeCKO cell collection included all three of the designed sgRNA sequences for the majority of the targeted loci in the pig genome (Fig. 1e). Finally, one of the originally designed sgRNA sequences (targeting the *IZUMO3* gene) was randomly selected to evaluate potential off-target effects (Supplementary Fig. 3a). A T7EN I cleavage assay revealed no off-target cleavage for any of the predicted potential off-target sites (Supplementary Fig. 3b). Collectively, this work demonstrates the development of a highly active and specific PigGeCKO resource with high utility for functional genomics research in pigs.

### JEV-induced cell death screening of the PigGeCKO cell collection to identify required host genes

We next developed a screening strategy, illustrated in Fig. 2a, to identify host genes required for successful JEV infection. To determine the optimal virus level for JEV-induced cell death in PK-15 cells for CRISPR screening, we examined JEV-induced cell death following infection at multiplicity of infection (MOI) of 0, 0.01, 0.05, and 0.1. As the infection dose of JEV was increased, we observed cytopathic effects (CPE) at approximately days 3-4 post virus infection; phenotypes included the rounding up and enlargement of cells, the formation of syncytia, and the detachment of cells into the medium (Supplementary Fig. 4a). Additionally, immunofluorescence analysis enabled us to detect the expression of the JEV-encoded NS3 protein in infected PK-15 cells (Supplementary Fig. 4b).

**Fig 2.**
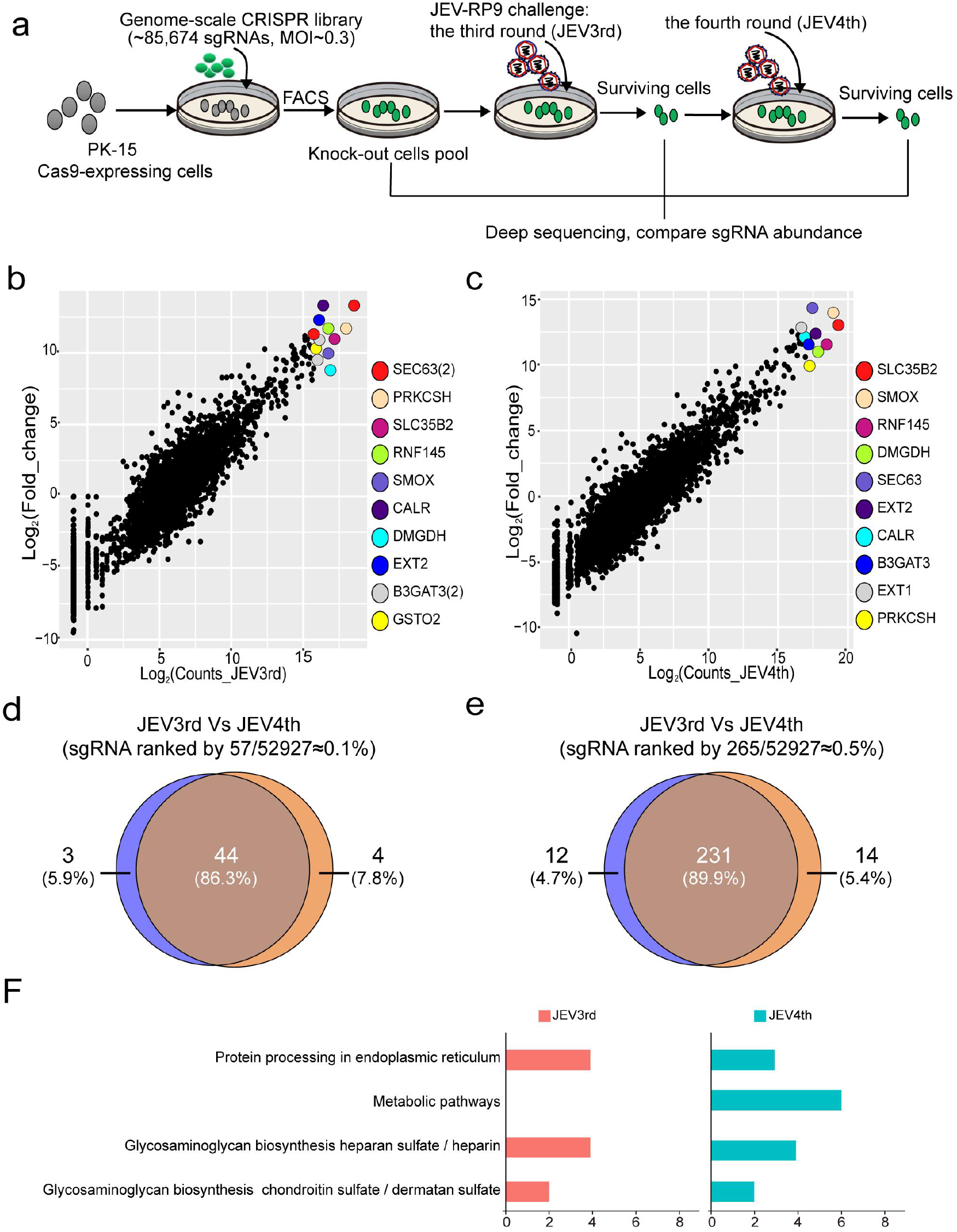
JEV resistance screen in PK-15 cells using the PigGeCKO cell collection. **a**, Workflow and screening strategy for the CRISPR/Cas9 screen. **b** and **c**, Scatter plots comparing sgRNA targeting sequences frequencies and extent of enrichment vs. the non-inoculated control mutant cell pool for the third (**b**) or fourth (**c**) rounds of JEV screens after challenge. **d** and **e**, Venn diagrams showing the overlapping enrichment of specific sgRNAs targeting sequences in the third or fourth rounds of JEV screens after challenge. For (**d**), among the top 0.1% of averaged reads for the sgRNAs; for (**e**), among the top 0.5%. **f**, KEGG pathway enrichment analyses for the top 0.5% ranked sgRNA targets from the third and fourth JEV challenge rounds. JEV: Japanese encephalitis virus; MOI: multiplicity of infection; sgRNA: small guide RNA; JEV3rd: the third challenge round of JEV screening; JEV4th: the fourth challenge round of JEV screening; FACS: Fluorescence-activated cell sorting; JEV-RP9: JEV, genotype 3, strain RP9.

Having established that PK-15 cells are a suitable model for identifying host genes with functions relating to JEV infection, we undertook our screen to identify host genes that modulate susceptibility to JEV-induced cell death. The PigGeCKO cell collection was infected with JEV and incubated for 11 days to enable selection of cells resistant to JEV-induced killing. We conducted four rounds of JEV challenge, employing untreated PK-15-Cas9 cells as a negative control to confirm the cell death caused by JEV infection in each round (Fig. 2a). At an MOI of 0.03, all untreated JEV-infected PK-15-Cas9 cells died, whereas a small number of viable cells from the JEV-infected PigGeCKO cell collection were detected. Surviving cells were collected and used for subsequent JEV challenge rounds, and the sgRNA constructs in surviving cells were PCR amplified and deep sequenced to identify candidate genes.

Focusing on protein coding genes, our screen found that a total of 2,181 unique sgRNA sequences were present in at least ten cells in the third and fourth rounds of JEV challenge. Among the originally designed 52,928 sgRNAs targeting 17,743 protein-coding genes (Supplementary Table 3), after JEV challenge, only 280 of the sgRNA constructs were present in at least 1,000 of the total population of analyzed cells (~0.5% of the total analyzed cells) (Supplementary Table 3). The top ten most enriched candidate genes after the third JEV challenge round were (highest-to-lowest) *SEC63*, *PRKCSH*, *SLC35B2*, *RNF145*, *SMOX*, *CALR*, *DMGDH*, *EXT2*, *B3GAT3*, and *GSTO2* (Figure 2B); the fourth challenge round identified the same enriched genes with the exception of *EXT1* for *SLC35B2* (Fig. 2c). When taking into consideration the design of three sgRNAs constructs for each targeted locus of the porcine genome, we found that multiple sgRNAs for the *SEC63*, *SLC35B2*, and *B3GAT3* genes were among the most highly enriched sequences after both the third and fourth rounds of JEV challenge (Fig. 2b, c).

After the third JEV challenge round, there were 57 sgRNA constructs present in at least 10,000 cells, and 219 sgRNA constructs present in at least 1,000 cells; after the fourth JEV challenge round, these numbers were, respectively, 57 and 239 (Supplementary Table 3). Comparison of enriched sgRNAs from the positive selection CRISPR screening revealed that 86.3% of the very highly enriched (i.e., ≥ 10,000) sequences were common to both the third and fourth challenge rounds and that 89.9% of the highly enriched (≥ 1,000) sequences were common to both rounds (Fig. 2d, e). These results highlight the capacity for CRISPR-based positive selection screening to consistently identify strong candidate genes. To explore the predicted biological functions of the candidate JEV-resistance genes, we performed KEGG pathway enrichment analyses for top 0.5% ranked sgRNA targets from the third and fourth JEV challenge rounds. These analyses revealed that the candidate JEV-resistance genes were significantly enriched for heparan sulfate proteoglycan metabolism and for Golgi and endoplasmic reticulum functions (Fig. 2f). Among these genes, *B3GAT3* and *EXT1* have known roles in JEV replication^28^. These results indicate that key host factors involved in JEV replication can be identified through multiple rounds of CRISPR screening.

### Knockout of Heparan sulfate proteoglycan (HSPG) pathway related genes significantly inhibits JEV entry

There are conflicting findings from previous studies of possible functional roles for heparan sulfate pathway proteins as cellular attachment factors during initiation of JEV infection^29,30^. HSPGs encompass a diverse class of proteins defined by the substitution with HS glycosaminoglycan (GAG) polysaccharide chains^31,32^. Our genome-scale CRISPR screen for JEV-infection related genes indicated that 10 genes associated with HSPG metabolism were among the most highly enriched sgRNA targeted genes: *EXT1*, *EXT2*, *GLCE*, *HS6ST1*, *B3GAT3*, *B4GALT7*, *XYLT7*, *EXTL3*, *SLC35B2*, and *GAA* (Fig. 3a). Among these genes, *SLC35B2*, *EXT1*, and *EXT2* were ranked in top 10 from both the third and fourth JEV challenge rounds. Notably, the *EXT1* and *HS6ST1* genes were each targeted by three separate sgRNA constructs, all of which were highly enriched, clearly indicating potential JEV-infection-related functions (Fig. 3b). HSPG synthesis and sulfation is driven by >20 different genes^28^; as shown in Fig. 3c and d, the significant enrichment of specific sgRNAs identified 7 genes potentially involved in HSPG synthesis and metabolic pathways, and 2 genes potentially involved in sulfurylation modifications of HSPG^33, 34^ in porcine cells.

**Fig 3.**
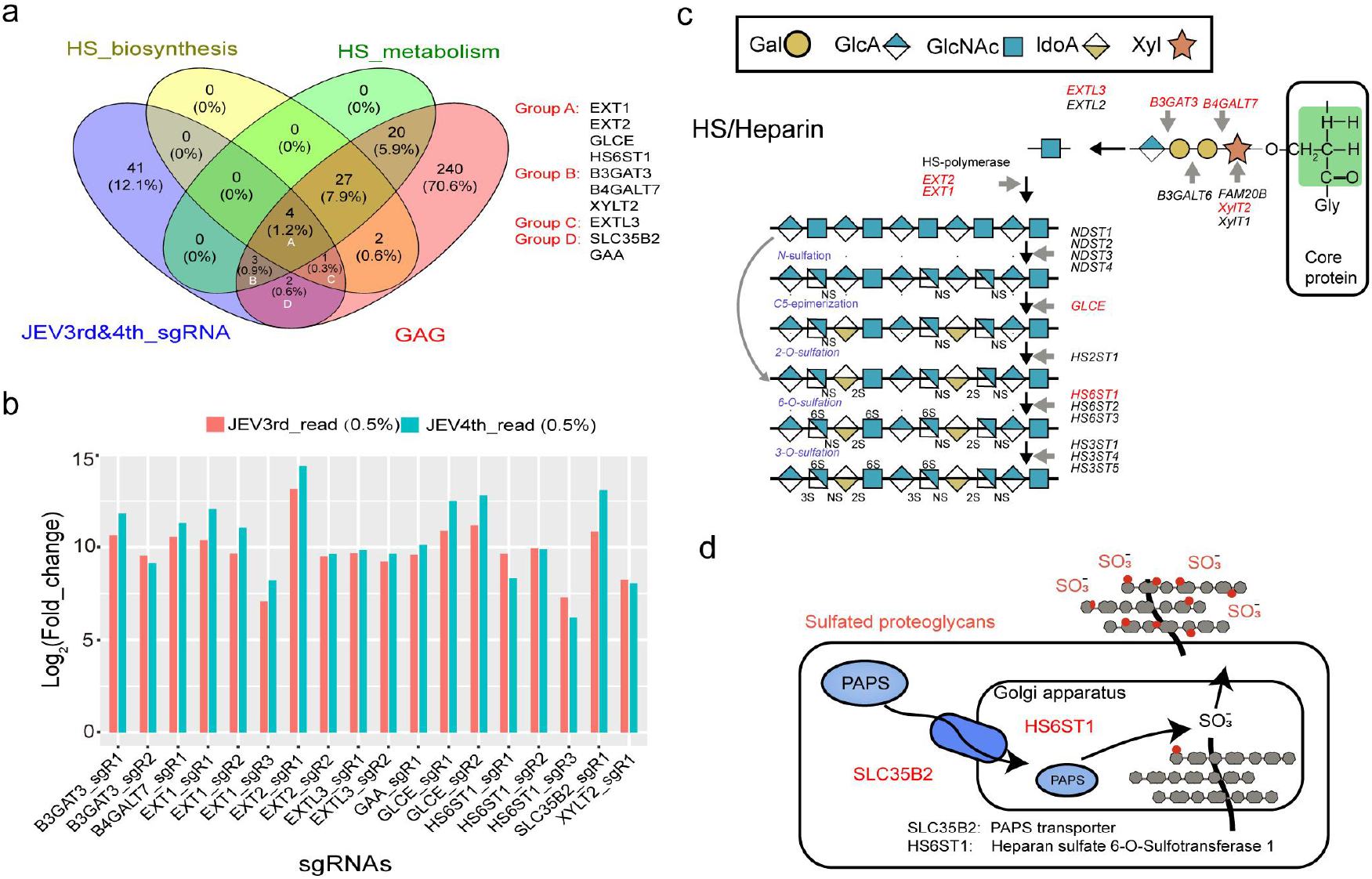
Significant enrichment of specific sgRNAs targeting 10 genes involved in HSPG synthesis and metabolic pathways. **a**, Venn diagram showing the overlapping enrichment of specific sgRNA targeted genes identified in the JEV screen of the cell knockout collection, HSPG biosynthesis and metabolism, and GAG metabolism pathways. **b**, Enrichment of specific sgRNAs targeting 10 genes involved in HSPG synthesis and metabolic pathways. **c**, Schematic diagram, adapted from Tanaka *et al*. (2017), showing the various classes of chemical components comprising HSPGs with known enzymes for their biosynthesis. Red indicated targeted genes found in this study. **d**, Schematic diagram, adapted from Blondel *et al*. (2016), showing the known components and localization information for the sulfurylation modifications known to occur for some HSPGs. Red indicated targeted genes found in this study. JEV: Japanese encephalitis virus; HS: Heparan sulfate; GAG: Glycosaminoglycan; HSPGs: heparan sulfate proteoglycans; PAPS: 3’-Phosphoadenosine-5’-phosphosulfate; JEV3rd: the third challenge round of JEV screening; JEV4th: the fourth challenge round of JEV screening.

Building from these initial candidate hits, we used CRISPR/Cas9 to generate *SLC35B2*, *HS6ST1*, *B3GAT3* and *GLCE* single knockout cell lines. Sanger sequencing confirmed that each of these monoclonal knockout cell lines had one or more nucleotide deletions predicted to cause a frameshift mutation in the coding regions of the targeted gene (a non-integer multiple of 3) (Fig. 4a). Viral loads in JEV-infected *SLC35B2*, *HS6ST1*, *B3GAT3*, and *GLCE* knockout cells and in wild-type PK-15 cells were measured at 18 hpi (hours post-infection) by plaque assay (Fig. 4b). In agreement with reduced viral loads observed in knockout cells, immunofluorescence assays showed that expression of the JEV-encoded NS3 protein in all four knockout cell lines was significantly reduced or undetectable following JEV infection at both 0.03 and 0.1 MOI (Fig. 4c). Next, cell cultures were sampled at 18 hpi for quantification of JEV genome copy number based on absolute quantitative PCR analysis using a pair of primers targeting the C gene of JEV. These analyses revealed that knockout of these four HSPG-related genes significantly inhibited JEV replication (12 hpi) (Fig. 4d). Furthermore, use of an antibody against Heparin/Heparan Sulfate antibody (10E4) to conduct immunofluorescence assays revealed that knockout of the *SLC35B2* and *HS6ST1* genes resulted in a significant reduction in the HSPG sulfurylation level compared to wild-type cells (Fig. 4e). This observation clearly suggests that sulfurylation modifications of HSPG can significantly and functionally impact the interaction between JEV and HSPG in PK-15 cells, potentially during viral entry. Finally, an EdU fluorescence assay showed no difference in cell proliferation rates of knockout and wide-type cells (Supplementary Fig. 5). Collectively, these results indicate that HSPG can act as a cellular adhesion factor or cofactor that mediates JEV entry.

**Fig 4.**
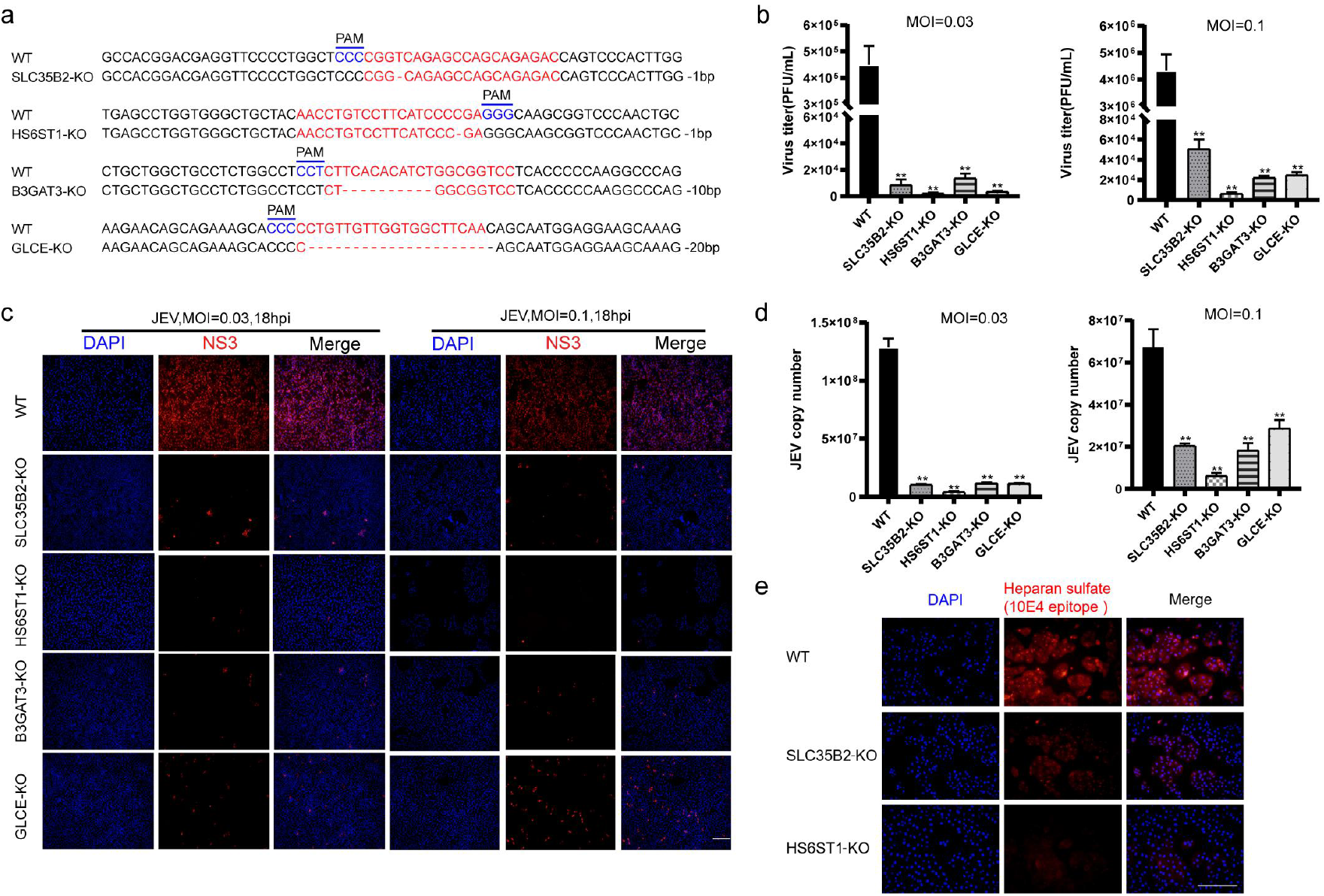
Knockout of genes coding for the HSPG pathway proteins SLC35B2, HS6ST1, B3GAT3, or GLCE significantly inhibits JEV replication in a dose-dependent manner in PK-15 cells. **a**, Sanger sequencing confirmation for the generated *SLC35B2*, *HS6ST1*, *B3GAT3*, and *GLCE* single gene knockout cell lines using CRISPR/Cas9 technology. **b**, Virus plaque assay for determination of viral concentration in SLC35B2, HS6ST1, B3GAT3, and GLCE knockout cells following infection with different JEV MOIs. **c**, Detection of JEV-encoded NS3 protein expressed in SLC35B2, HS6ST1, B3GAT3, and GLCE knockout cells (under different JEV MOIs) by immunofluorescence. **d,** Determination of JEV copy number in SLC35B2, HS6ST1, B3GAT3, and GLCE knockout cells (under different JEV MOIs) based on absolute quantitative real-time PCR. **e**, Detection of HSPG sulfurylation level in SLC35B2 and HS6ST1 knockout cells by immunofluorescence. PAM: protospacer adjacent motif; JEV: Japanese encephalitis virus; MOI: multiplicities of infection; hpi: hours post-infection; DAPI: 4’,6-diamidino-2-phenylindole; KO: knockout cells; WT: wild-type cells. * p ≤ 0.05; ** p ≤ 0.01; Scale bar, 100 μm.

### EMC3 is required for JEV replication

The endoplasmic reticulum membrane complex (EMC) is known to be required for infection by flaviviruses, which have RNA genomes^35,36^. However, it is unclear whether EMC family genes are involved in JEV replication. Interestingly, our genome-scale CRISPR JEV infection screen showed that *EMC3* and *EMC6* genes, both of which encode ER membrane protein complex subunits, were ranked 44 and 34 among the candidate hits in the fourth JEV challenge round, respectively (Supplementary Table 3). As such, we generated two independent *EMC3* null cell lines using CRISPR/Cas9. Sanger sequencing confirmed the presence of 1 or 2 bp deletions or insertions (a non-integer multiple of 3) in both *EMC3* knockout cell lines (Fig. 5a), and immunoblotting confirmed that the EMC3 protein was not expressed in cells of either knockout line (Fig. 5b).

**Fig 5.**
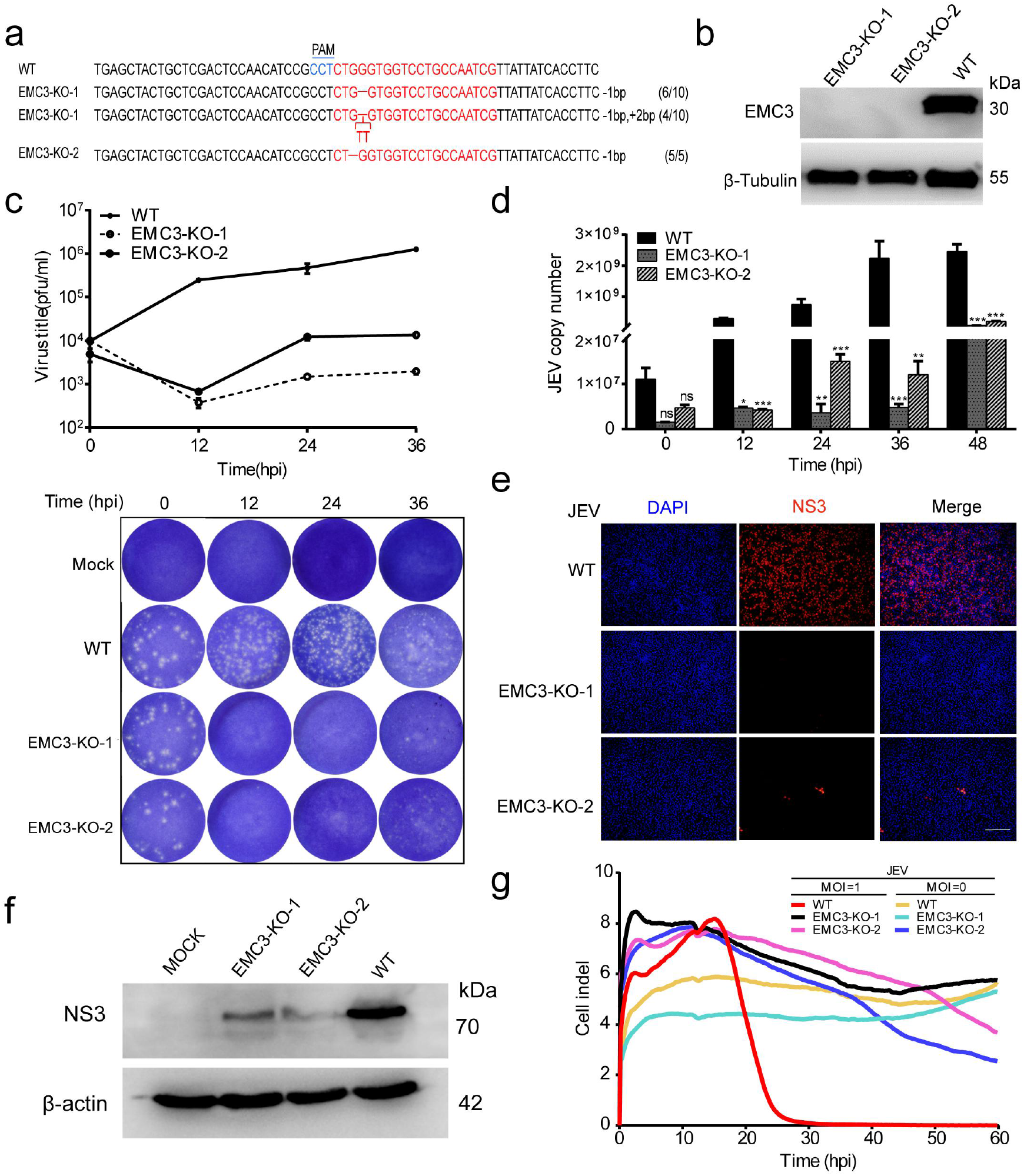
The endoplasmic reticulum membrane protein complex subunit 3 (EMC3) is required for JEV replication. **a**, Sanger sequencing confirmation for generation of two EMC3 knockout cell lines (EMC3-KO-1 and 2) using CRISPR/Cas9 technology. An example target region with CRISPR target sites highlighted in red, and the PAM high-lighted in blue. **b**, Detection of EMC3 protein by immunoblotting. **c**, Determination of viral concentration in EMC3 knockout cells by virus plaque assay. **d**, Determination of JEV copy number in EMC3 knockout cells by absolute quantitative real-time PCR. **e**, Detection of JEV-encoded NS3 protein expressed in EMC3 knockout cells by immunofluorescence. **f**, Detection of JEV-encoded NS3 protein expressed in EMC3 knockout cells by immunoblotting. **g**, Comparison of the ability of EMC3 knockout and wild-type cells to resist JEV-induced cell death (MOI = 1) using a Real-Time Cell Analyzer (RTCA). PAM: protospacer adjacent motif; JEV: Japanese encephalitis virus; MOCK: non-infected cells was included as negative control; MOI: multiplicities of infection; hpi: hours post-infection; DAPI: 4’,6-diamidino-2-phenylindole; KO: knockout cells; WT: wild-type cells; kDa: kilodalton. * p ≤ 0.05; ** p ≤ 0.01; ***p ≤ 0.001; Scale bar, 100 μm.

Subsequently, viral concentrations in JEV-infected *EMC3* null and wild-type PK-15 cells were determined at 0, 12, 24, and 36 hpi by both plaque assay and quantification of JEV genome copy number based on absolute quantitative PCR analysis using a pair of primers targeting the C gene of JEV. Together, these analyses revealed that knockout of the *EMC3* gene significantly inhibited JEV replication (12 hpi) (Fig. 5c, d). Both JEV-infected EMC3 knockout cell lines possessed substantially reduced levels of viral NS3 protein expression as determined via immunofluorescence analysis (Fig 5E) and immunoblotting (Fig 5F). Finally, the ability of *EMC3* knockout cells to resist JEV-induced death at a high dose (MOI = 1) was evaluated. The result from cultures grown using a Real-Time Cell Analyzer (RTCA) verified that *EMC3* knockout cells were able to completely resist JEV-induced cell death (Fig. 5g). Collectively, these results demonstrate that the EMC3 is required for JEV-induced PK-15 cell death.

### CALR is required for JEV replication

Among the top 10 ranked genes in the genome-scale CRISPR screen for JEV infection screening candidates was a gene known to function in intracellular calcium homeostasis: *CALR*. To explore the potential function of CALR in mediating JEV replication, *CALR* knockout cells was generated by CRISPR/Cas9. Sanger sequencing showed that the selected monoclonal *CLAR* knockout cells have a 1 bp insertion (Fig. 6a), and immunoblotting confirmed that the CALR protein was not expressed in the knockout cells (Fig. 6b).

**Fig 6.**
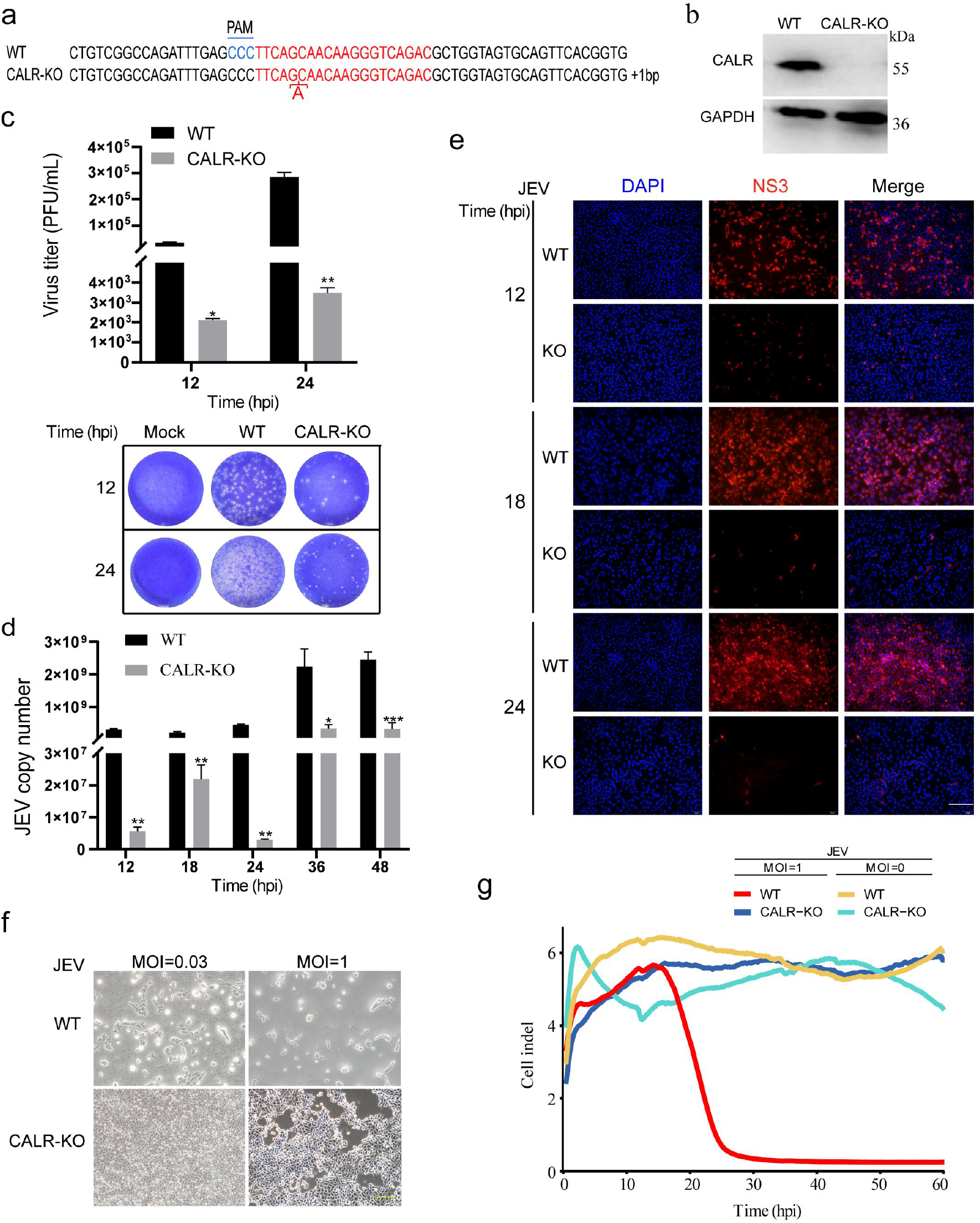
The calreticulin (CALR) calcium-binding protein of the endoplasmic reticulum lumen is required for JEV replication. **a**, Sanger sequencing confirmation for generation of a CALR knockout cell line using CRISPR/Cas9 technology. An example target region with CRISPR target sites highlighted in red, and the PAM high-lighted in Blue. **b**, Detection of CALR protein in knockout cells by immunoblotting. **c**, Determination of viral concentration in CALR knockout cells through a virus plaque assay. **d**, Determination of JEV copy number in CALR knockout cells by absolute quantitative real-time PCR. **e**, Detection of JEV-encoded NS3 protein expressed in CALR knockout cells by immunofluorescence. **f**, Comparison of the ability of CALR knockout and wild-type cells to resist JEV-induced cell death using cell proliferation assays under various JEV MOIs. **g**, Comparison of the ability of CALR knockout and wild-type cells to resist JEV-induced cell death (MOI = 1) using a Real-Time Cell Analyzer (RTCA). PAM: protospacer adjacent motif; JEV: Japanese encephalitis virus; MOCK: non-infected cells was included as negative control; MOI: multiplicities of infection; hpi: hours post-infection; DAPI: 4’,6-diamidino-2-phenylindole; KO: knockout cells; WT: wild-type cells; kDa: kilodalton. * p ≤ 0.05; ** p ≤ 0.01; ***p ≤ 0.001; Scale bar, 100 μm.

Next, plaque assays and absolute quantitative real-time PCR were used to measure viral concentrations in JEV-infected *CALR* null and wild-type PK-15 cells at 12 and 24 hpi. Concurrently, JEV-infected cell cultures were harvested at 12, 18, 24, 36 and 48 hpi, and viral RNA was extracted from cell suspensions and cDNAs were synthesized as absolute quantitative PCR template. As shown in Fig. 6c and d, knockout of the *CALR* gene significantly inhibited JEV replication. Moreover, immunofluorescence results showed that the NS3 protein was only weakly expressed at 12, 18 and 24 hpi in JEV-infected *CALR* null cell lines (Fig. 6e). Finally, the ability of *CALR* knockout cells to resist JEV-induced death at a high challenge dose (MOI = 1) was evaluated. Results of the cell proliferation assay and Real-Time Cell Analyzer assay showed that, compared to wild-type cells, knockout of *CALR* can confer resistance to JEV-induced death (Fig. 6f, g). Furthermore, in the EdU fluorescence assay, the number of fluorescent cells was consistent in *CALR* knockout cells and wild-type cells, indicating that knocking out CALR genes did not affect normal cell proliferation (Supplementary Fig. 6). These results indicate that CALR is required for JEV replication.

## DISCUSSION

Our results highlight the power of CRISPR/Cas9-based screening for functional analyses in pigs, and we present in Figure 7 a preliminary proposed model for JEV entry and replication in PK-15 cells based on our findings. The CRISPR screens recovered JEV entry cofactor HSPG-related host factors (SLC35B2, HS6ST1, EXT1, EXT2, GLCE, B3GAT3, B4GALT7, XYLT7 and EXTL3), as well as multiple host factors involved in calcium homeostasis (CALR), and transmembrane protein processing and maturation (EMC3 and EMC6). In addition, a large number of other candidate host factors involved in JEV infection of host cells were identified with our CRISPR screen, warranting further investigation.

**Fig 7.**
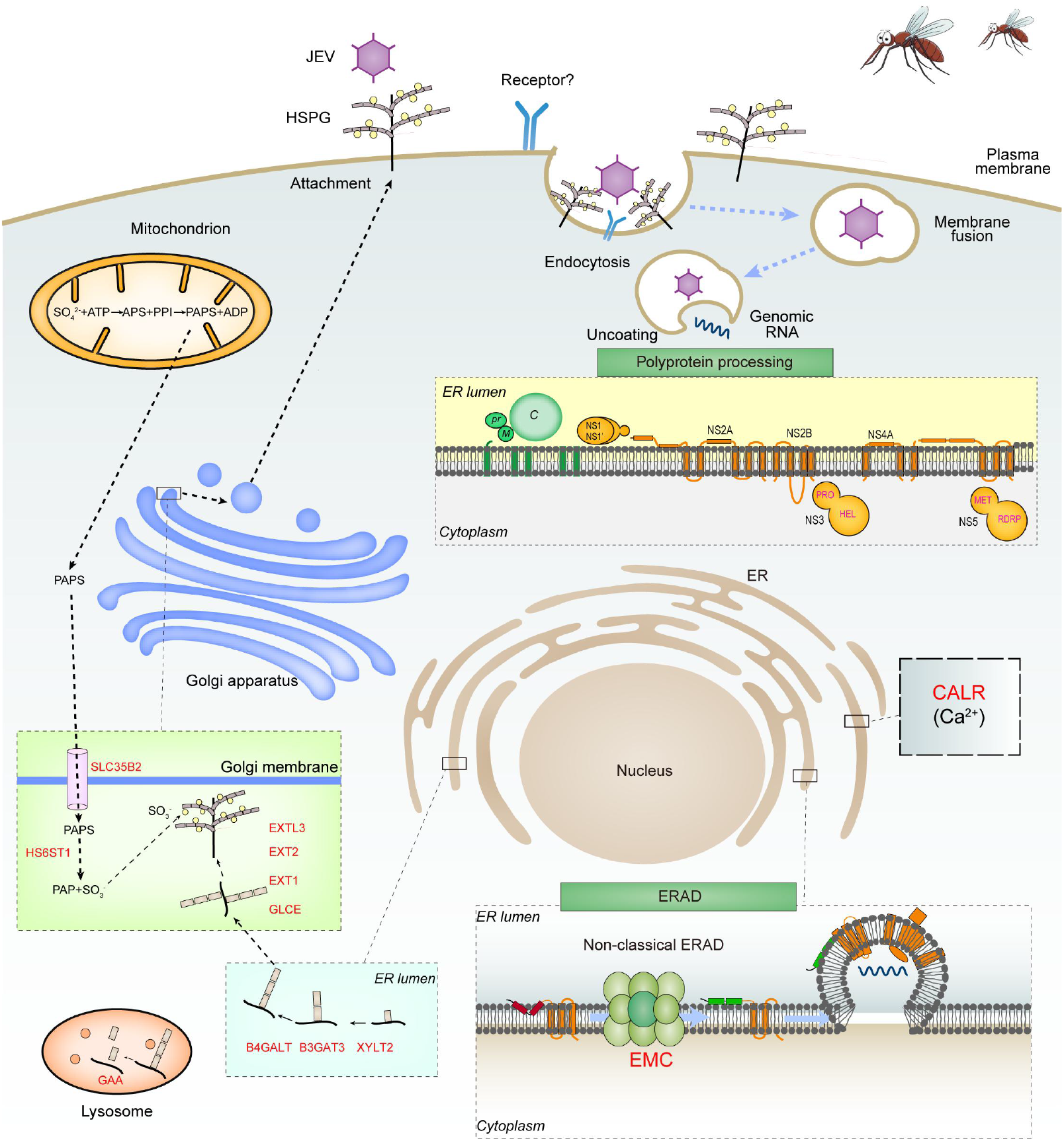
A proposed model for JEV entry and replication in PK-15 cells. The JEV infection cycle starts with binding to co-factors Heparan sulfate proteoglycan (HSPG), and/or unknown cellular receptors, followed by viral entry to enable replication. The endoplasmic reticulum membrane protein complex (EMC) and calreticulin (CALR) calcium-binding protein of the endoplasmic reticulum (ER) lumen are involved in JEV replication of host cells. Subsequently, the JEV RNA genome is replicated, viral particles are matured and packaged, and are released from cells. JEV: Japanese encephalitis virus; HSPG: heparan sulfate proteoglycans; PAPS: 3’-Phosphoadenosine-5’-phosphosulfate; PAP: 3’-phosphoadenosine 5’-phosphate; ATP: Adenosine triphosphate; ADP: Adenosine diphosphate; APS: Adenosine 5’ phosphosulfate. PPI: pyrophosphate; ER: endoplasmic reticulum; ERAD: endoplasmic reticulum-associated protein degradation; EMC: endoplasmic reticulum membrane protein complex.

While genome-scale CRISPR/Cas9 mutagenesis methods obviously facilitate gene functional studies in both cellular and animal models^19–24^, and the ability to programmably target the entire coding or regulatory genome represents a significant advance over spontaneous or random mutagenesis, genome-scale CRISPR/Cas9 approaches share with classical forward genetic screens the requirement for an assay to enrich for cells exhibiting the phenotype of interest^37–39^. JEV causes encephalitis in humans and reproductive disorders in pigs, the latter leading to substantial economic losses^10^. In the present study, we developed a series of genome-scale pig CRISPR/Cas9 knockout library (PigGeCKO) resources that facilitate the pooled screening of genes that prevent JEV invasion or replication, thereby inhibiting JEV-induced cell death in porcine cells. Furthermore, we demonstrated how lentiviral delivery of a PigGeCKO library targeting 17,743 protein-coding genes enables positive selection screening for JEV-replication associated host genes.

The first candidate genes examined from this screen are known to function in HSPG metabolism, specifically in the synthesis and modification of heparan sulfate chains in normal cells. The murine homolog of EXT1 protein is known to be localized to the Golgi apparatus^40^, where it binds with EXT2 to form a complex known to modify heparan sulfate^40^. EXT1, EXT2, and EXTL3 together contribute to heparan sulfate chain elongation^41^. In mice, GLCE is an epimerase enzyme required for the biosynthesis of HSPGs, which are composed of a core protein and one or more heparan sulfate glycosaminoglycan chains^42^. The B3GAT3 enzyme catalyzes the formation of glycosaminoglycan-protein linkages by glucuronic acid transfer, which is the final step in the biosynthesis of proteoglycan-linked regions^28^. A previous study identified that knocking out the *B3GAT3* gene can significantly inhibit JEV replication^28^, a conclusion that was further confirmed by our genome-scale CRISPR screening and subsequent hypothesis-driven functional analyses. A different study showed that the reduced activity of the B4GALT7 enzyme is associated with a reduced substitution of the proteoglycans decorin and biglycan (both of which have glycosaminoglycan carbohydrate chains) and alterations in heparan sulfate biosynthesis^43^.

Our work strongly supports that HSPG pathway genes can mediate JEV replication in PK-15 cells. Previous studies have shown that the *SLC35B2* gene product is located in the microsomal membrane and functions to transport 3’-Phosphoadenosine-5’-phosphosulfate (PAPS) from the cytosol (where it is synthesized) into the Golgi lumen^43^, whereas the HS6ST1 sulfotransferase enzyme uses PAPS as a substrate for heparan sulfate biosynthesis^44,45^. Accordingly, our immunofluorescence assays showed that *SLC35B2* or *HS6ST1* gene knockout in PK-15 cells significantly reduced the extent of sulfurylation modification to heparan sulfate. Therefore, we believe that sulfurylated HSPG serves as an adjunct or adhesion factor during JEV invasion of host cells. It should be noted that our initial screening efforts were based on relatively low MOI JEV challenges of PK-15 mutant cells. Although these experiments succeeded in identifying many host factors involved in JEV replication, future CRISPR screening efforts employing a higher viral challenge dose would potentially reduce false positives and/or identify additional critical host factors in this process.

The second host cellular process implicated in JEV infection involves the EMC3 subunit of the ER membrane protein complex, which is involved in ER-mitochondrial membrane tethering and which is required to facilitate lipid transfer from the ER to the mitochondrial membrane, impacting nearly all aspects of cell physiology^46^. Previous research has found that the ER membrane protein complex is a transmembrane domain insertase, thus loss of EMC causes ER stress and altered protein trafficking^47^. For ZIKV, this complex was required for viral protein accumulation in a cell line harboring a ZIKV replicon^48^. Our CRISPR screen showed that knockout of *EMC3* and *EMC6* can inhibit JEV-induced cell death. In particular, knockout of the *EMC3* gene significantly inhibited the replication of JEV in PK-15 cells. EMC3 and EMC6 are associated with the ERAD pathway, which may participate in the secretory protein quality control processes that guide the removal of aberrantly folded proteins from the ER. Thus, whether the EMC subunits participate in JEV protein biogenesis, misfolding, or direct interaction with JEV particles or JEV-encoded proteins requires further study, for example by using small molecule inhibitors of ER stress.

The final gene we examined via follow-up hypothesis driven studies was *CALR*, which encodes a multifunctional soluble protein that can bind Ca^2+^ ions. Knockout of *CALR* resulted in a strong JEV resistance phenotype. Proper folding in the endoplasmic reticulum is a prerequisite for the correct localization and function of most secreted and transmembrane proteins^49^, and previous studies have found that *CALR* and *calnexin* (CANX) chaperones mediate nascent glycoprotein folding in the endoplasmic reticulum^50,51^. Thus, we hypothesize that *CALR* appears to be an essential gene that links JEV replication to downstream cell death pathways, possibly due to calcium homeostasis disequilibrium. For these reasons, *CALR* knockout cells represent a highly useful new model for studying the relationship between calcium ion homeostasis and JEV infection, additional studies in this area are currently ongoing.

While this study validated six genes selected with follow-up knockout studies and infection assays, our positive selection screening strategy yielded many candidates that may function in JEV infection and as such merit further investigation. We plan to extend the screening described herein to achieve full genome saturation to increase the search scope for host factor genes to further deepen our understanding of the multifaceted and spatiotemporally programmed interactions between JEV and host cells. One limitation of this study is that it did not specifically address issues of potential cell specificity of JEV infection. Although PK-15 cells are susceptible to JEV, this virus primarily infects nerve cells and germ cells^10,25^. It is also important to note that JEV can infect many different types of cells from highly divergent species, including but not limited to Vero cells, BHK-21 cells, and mouse neuro 2a cells^14^. Thus, testing whether the required factors we identified for JEV infection of PK-15 cells play the same roles in other cell types will require further functional validation.

It is important to consider that some genes may be necessary for normal cell growth, so their knockout may inhibit cell proliferation, which would prevent detection of required genes. JEV is an RNA virus and belongs to the flaviviridae family, and previous studies in humans using CRISPR-based screening strategies have identified many required host genes for cell death induced by DENV, ZIKV, WNV, YFV, and HCV; it is notable that few of these genes appear to overlap^20–24^. On one hand, the identified genes may differ owing to heterogeneities in the various cell types or species; on the other hand, perhaps the host factors for JEV infection simply differ greatly from other flaviviridae family viruses. Importantly, besides the protein-coding genes examined here, our CRISPR screening additionally identified candidate JEV-replication related lncRNAs (data not shown). Our results highlight the complexity of JEV entry, replication, packaging, and release from host cells, yet also lead the way for a variety of hypothesis-driven basic biological and medical studies to deepen our understanding of this complex process.

## MATERIALS and METHODS

### Plasmids

The lenti-Cas9-Puro and lenti-sgRNA-EGFP vectors were kindly provided by Professor Xingxu Huang at ShanghaiTech University. The lenti-Cas9-Puro vector was used for generation of the Cas9-expression cell line (PK-15-Cas9). To construct the lentiviral sgRNA vector, paired oligonucleotides of sgRNA (50 μM per oligo) were annealed and cloned into lenti-sgRNA-EGFP which was linearized with *Bbs*I (Supplementary Table 4). All plasmids were confirmed by Sanger sequencing (Tsingke).

### Genome-wide porcine sgRNA library design

Three sgRNAs were designed against each protein-coding gene, lncRNA, and miRNA using software of CRISPR-offinder (version 1.2, http://www.biootools.com^26^). Sequences of protein-coding genes, lncRNA, and miRNA, were found from databases of Ensemble (version 10.2, www.ensembl.org/index.html), ALDB (http://202.200.112.245/aldb^52^), and miRBase (www.mirbase.org^53^), respectively. Briefly, the selected sgRNAs were weighted based on targeting the first 50% of the open reading frames and minimizing potential off-target sites. The maximum number of mismatches allowed up to 3 nucleotides to the DNA target in the 20 mer targeting region of selected sgRNAs, targeting the miRNA hairpin region, and to avoid overlap between sgRNAs in the same given targets.

### Construction of a genome-wide sgRNA library plasmid

The sgRNA library was synthesized using CustomArray 90K arrays (CustomArray Inc.), and amplified by PCR using Phusion High-Fidelity PCR Master Mix with HF Buffer (NEB) to produce sub-pools for Gibson assembly (NEB). The PCR reaction was performed in a Veriti™ 96-Well Thermal Cycler (Thermo Fisher Scientific) with 16 cycles. In total, 40 PCR reactions were performed using 50 ng of oligo pool per 50 μL of reaction volume. The PCR products were mixed and purified using a MinElute PCR purification Kit (QIAGEN), and then ligated into the linearized lenti-sgRNA-EGFP vector using Gibson assembly. The ligation mixture were transformed into Trans1-T1 Phage Resistant Chemically Competent Cells (Transgen). To achieve sufficient coverage, parallel transformations were performed, counting the number of colonies to reach 200-times total sgRNA in the library. The sgRNA library plasmids were extracted with a Plasmid Plus Maxi Kit (QIAGEN). The library plasmids were amplified using PrimeSTAR GXL DNA Polymerase (Takara) with 16 reaction cycles. PCR products were purified using a QIAquick Gel Extraction Kit (QIAGEN) and then analyzed by high-throughput sequencing to examine the sgRNA coverage in the library plasmids. All primers for constructing sgRNA expression vector are listed in Supplementary Table 4.

### Cell culture

PK-15, HEK293T, and BHK-21 cell lines were purchased from the Cell Bank of the Chinese Academy of Sciences (Shanghai, China), and were subjected to mycoplasma detection. For all experiments, cells were maintained in Dulbecco’s Modified Eagle Media (DMEM) supplementing with 10% fetal bovine serum (FBS) and 100 U/mL penicillin and 100 μg/mL streptomycin and incubated at 37°C with 5% CO_2_.

### Generation of Cas9-expression cell line

PK-15 cells were transduced with Cas9-puro lentivirus. At 3 days after transduction, cells were changed to fresh growth medium containing 3 μg/mL puromycin. Antibiotics resistant cells were collected and reseeded into 100 mm dishes at a concentration of 100 cells per dish to generate single-cell clones. At 7 days after antibiotics selection, the single-cell clones were tested for Cas9 expression by immunoblotting. In addition, the *ANPEP* gene targeted by a sgRNA was used to screen cell lines with high expression level and activity of Cas9. The resulting cell lines were designated as PK-15-Cas9.

### sgRNA library lentivirus production and transduction

To produce lentivirus, co-transfection of 12 μg of the library plasmid, 4 μg of pMD2.G plasmid (Addgene), and 8 μg of psPAX2 (Addgene) plasmid per 100 mm dish by using JetPRIME (PolyPlus) according to the manufacturer’s instructions. At 60 hrs post-transfection, the cell supernatants were collected, filtered by using a 0.45 μm low protein binding membrane (Millipore), and then centrifuged at 30,000 rpm and 4°C for 2.5 hrs. The virus pellets were resuspended in phosphate buffered saline (PBS, pH7.4), aliquoted and stored at −80°C. Target cells were transduced with the resulting lentiviruses in the presence of 8 μg/mL polybrene (Sigma-Aldrich). At 24 hrs after transduction, viruses were removed and replaced with fresh media.

### Generation of mutant cell libraries and screening

A total of ~2×10^8^ PK-15-Cas9 cells were seeded into T225 flasks and infected with the library lentiviruses at multiplicity of infection (MOI) of 0.3. Three days post-infection, GFP-positive cells were collected by FACS (Fluorescence-Activated Cell Sorting) and reseeded into 100 mm dishes. Six days post-infection, DNA from ~7×10^6^ cells was extracted using a Blood & Cell Culture DNA Midi Kit (QIAGEN) and amplified to examine the coverage of mutant cell libraries. For the CRISPR screening, ~6×10^7^ mutant cells were infected with JEV-RP9 at an MOI of 0.03 in DMEM without FBS and incubated at 37°C and 5% CO_2_. After 1.5 h incubation, the inoculum was removed and replaced with fresh DMEM supplementing with 2% FBS and 1% penicillin-streptomycin. At 11 days post-infection, viable cells were collected and expanded for the deep sequencing analysis and the next round of infection.

### Knockout of candidate gene in PK-15 cells by CRISPR/Cas9

Individual sgRNA targeting to candidate gene was cloned into the linearized lenti-sgRNA-EGFP and lentivirus were produced as described above. The resulting lentivirus was transduced into PK-15-Cas9 cells. Cells with GFP expression were enriched by FACS and then seeded into 96-well plates to generate single-cell clones. 7 days after transduction, the genotypes of cell colonies were analyzed by extracting genomic DNA (TIANamp Genomic DNA Kit, TIANGEN) and sequencing. All primers for identifying the genotype of cell colonies are listed in Supplementary Table 4.

### T7 endonuclease I cleavage detection assay and Sanger sequencing

All potential off-target sites with high homology in the sgRNAs were predicted using software CRISPR-offinder^26^. Genomic DNAs were extracted using the TIANamp Genomic DNA Kit (TIANGEN) from mutated clonal cells for PCR amplification, and T7 endonuclease I (T7EN I) cleavage detection assay was employed to determine off-target effects. CRISPR/Cas9-induced lesions at the endogenous target site and predicted off-targets were quantified using the T7EN I cleavage detection assay to investigate the insertions/deletions (indels) generated by nuclease-mediated non-homologous end joining (NHEJ). The gene fragments of off-target sites were amplified with primers specific to each locus by 35 cycles of PCR with TaKaRa LA Taq (TaKaRa). The PCR products were purified and subjected to the process of denatured and annealed by using a thermocycler, and the hybridized PCR products were digested with T7EN I (NEB) for 15 min and separated with a 2% agarose gel. The agarose gels were stained with Gel-Red and the signal of DNA in the gel was quantified by densitometry using ImageLab software (Bio-Rad). All primers are listed in Supplementary Table 4.

### Illumina sequencing of sgRNAs in the genome-wide library and enriched mutants

The genomic DNA of each sample was extracted using a Blood & Cell Culture DNA Midi Kit (QIAGEN). The sgRNA-coding region was amplified by PCR using Q5^®^ Hot Start High-Fidelity DNA Polymerase (NEB) in a reaction volume of 50 μL. PCR products were mixed and purified with a MinElute PCR purification Kit (QIAGEN). The purified PCR products were amplified by PCR using different barcoded primers. All PCR products were pooled and purified with a MinElute PCR purification Kit (QIAGEN), followed by Illumina Next-generation sequencing. Mapped read counts were subsequently used as input for the MAGeCK analysis software package (version 0.5)^54^. Then, the top 0.5% ranked sgRNAs from the third and fourth JEV challenge rounds were used to identify enriched targeting protein-coding genes. Kyoto Encyclopedia of Genes and Genomes (KEGG) enrichment analyses were performed in the Database for Annotation, Visualization and Integrated Discovery (DAVID) (https://david.ncifcrf.gov/)^55^. All primers are listed in Supplementary Table 4.

### Virus plaque assay

Plaque assays were performed on BHK-21 cells. Briefly, BHK-21 cell monolayers at 50% confluence were incubated with serially diluted virus at 37°C with 5% CO_2_. 2 hrs after incubation, inoculum was removed, and the cells were overlaid with 50% 2× DMEM, 50% Agarose LMP (Genview), 2% FBS and 1% penicillin streptomycin for 3 days. Cells were fixed with 10% formaldehyde neutral solution overnight at room temperature, then stained with 0.5% crystal violet for 2 hrs at room temperature. Plaques were counted manually and plaque-forming units were calculated. Three independent experiments were performed, with results presented as mean ± SD.

### Absolute quantitative real-time PCR

Viral RNA was extracted from cell suspensions using a Viral RNA Extraction Kit (TaKaRa) and following the manufacturer’s protocol. About 1 μL of viral RNAs were used as template to synthesize cDNAs with PrimeScript™ RT reagent Kit with gDNA Eraser (TaKaRa). Absolute quantitative real-time PCR assay was performed with SYBR green I (TOYOBO) and primers binding to C gene of JEV in a final reaction volume of 20 μL. Primers used in quantitative PCR are listed in Supplementary Table 4.

### Immunofluorescence assay

The expression level of JEV NS3 protein in wild-type or gene knockout PK-15 cells as indicated in figures was determined by immunofluorescence assay. Briefly, cells grown on the glass coverslip in 6-well cell culture plates were infected with JEV at different MOI. After infection with different setting times, cells were fixed with 4% paraformaldehyde for 15 minutes at room temperature, washed twice with pre-cooled PBS, and then permeabilized for 10 minutes at room temperature with cold 0.3% TritonX-100 in PBS. Cells were reacted with NS3 (JEV) antibody (GTX125868, GeneTex) or anti-Heparin/Heparan Sulfate antibody (10E4, US Biological) at 4°C overnight, and the primary antibodies were recognized by Alexa Fluor^®^ 594 Donkey anti-rabbit IgG (H+L) (ANT030, Antgene) or Alexa Fluor^®^ 555 Anti-mouse IgG (H+L) (4409S, Cell Signaling Technology) and shaken at room temperature for 1.5 hrs in the dark. Cell nuclei were counter-stained with DAPI (4’, 6-diamidino-2-phenylindole) for 10 minutes at room temperature in the dark. Cells were observed with a fluorescence microscope (OLYMPUS IX3-RFACS), and images were taken with OLYMPUS DP80 camera.

### Immunoblotting assay

Approximately 1.2×10^6^ of PK-15 cells were lysed in ice-cold cell lysis buffer (50 mM Tris-HCl [pH6.8], 2% SDS, 10% glycerol, 1% β-mercaptoethanol, 12.5 mM EDTA, 0.02% bromophenol blue) supplemented with protease inhibitor (Beyotime) and phenylmethylsulfonyl fluoride (PMSF) (Beyotime), with cell lysates precleared at 4°C by centrifugation at 13, 000 rpm for 10 minutes. The precleared lysates were separated by 10% polyacrylamide gel SDS. Separated proteins were then transferred onto a nitrocellulose membrane and probed with Cas9 (GTX53807, GeneTex), EMC3 (sc-99670, Santa Cruz Biotechnology), CALR (A1066, Abclonal) antibody, with β-tubulin (Sungenebiotech)/GAPDH antibodies (Beyotime) used as an internal loading control. The primary antibodies were detected with horseradish peroxidase (HRP) conjugated goat anti-rabbit IgG (AS014, Abclonal) or goat anti-mouse IgG (A0208, Beyotime), and the secondary antibodies were visualized by ECL Prime Western Blotting Detection Reagents (GE Healthcare, UK).

### Statistical analysis

Statistical analysis was performed using R programming language. The mean ± SEM was determined for each treatment group in the separated experiments. Two-tailed Student’s t-test was used to determine significant differences between treatment and control groups (*P* ≤ 0.05).

## Supplemental information

Supplemental Information includes 6 figures and 4 tables and can be found with this article online.

## Author contributions

Most of the experimental work was co-conducted by CZ Zhao, HL Liu, and TH Xiao, with minor contributions from ZC Wang, XS Han, JF Zhang, LX Qin, JX Ruan, MJ Zhu YL Miao and B Zuo. XW Nie helped in analyzing the data and drawing figures, P Qian provided the JEV-PR9 strain and technical support. SS Xie conceived the project, designed the experiments and wrote the manuscript. K Yang and XY Li helped in revising the manuscript. SH Zhao provided support and supervised the project. All authors contributed to manuscript revision.

## Acknowledgement

This work was supported by the National Transgenic Project of China (grant numbers: 2018ZX08009-26B and 2016ZX08006003-004), the Fundamental Research Funds for the Central Universities and the Natural Science Foundation of China, and the National Key Research and Development Program of China, Stem Cell and Translational Research (2016YFA0100203). We thank Dr. Erwei Zuo for critical review of the manuscript. We thank Thuy-Nhien Tran-Thi for linguistic assistance during the preparation of this manuscript. We also thank Dr. Yan Wang (Institute of Hydrobiology, Chinese Academy of Sciences) for her assistance with flow cytometry.

